# Sustained TNF signaling is required for the synaptic and behavioral response to acute stress

**DOI:** 10.1101/2021.12.22.473829

**Authors:** Gina M Kemp, Haider F Altimimi, Yoonmi Nho, Renu Heir, David Stellwagen

## Abstract

Acute stress triggers plasticity of forebrain synapses as well as behavioral changes. Here we reveal that Tumor Necrosis Factor α (TNF) is a required downstream mediator of the stress response in mice, necessary for stress-induced synaptic potentiation in the ventral hippocampus and for an increase in anxiety-like behaviour. Acute stress is sufficient to activate microglia, triggering the long-term release TNF. Critically, on-going TNF signaling in the ventral hippocampus is necessary to sustain both the stress-induced synaptic and behavioral changes, as these could be reversed hours after induction by antagonizing TNF signaling. This demonstrates that TNF maintains the synaptic and behavioral stress response *in vivo*, making TNF a potential novel therapeutic target for stress disorders.

## Introduction

Stress drives adaptive responses critical for an organism’s physiological homeostasis and function. However, unregulated activation of the stress response results in significant problems; stress is a major contributing factor to the development of a number of psychiatric diseases including posttraumatic stress disorder and other anxiety disorders (Brewin, 2005; McEwen et al., 2015). There are long-standing observations of reciprocal interactions between stress and the immune system; stimulation of the immune system is a potent activator of the hypothalamic-pituitary-adrenal (HPA) axis, and the output of the HPA axis (corticosteroid (Cort)) can exacerbate inflammation in the brain under certain conditions or be an immune suppressant in other conditions (Costa-Pinto and Palermo-Neto, 2010; Dhabhar, 2009; Sorrells et al., 2009). Further, elevated pro-inflammatory cytokine levels are observed in patients with generalized anxiety disorder (Hou et al., 2017; Tang et al., 2018) and even acute stressors can increase cytokines levels in humans (Muscatell et al., 2015; Slavish et al., 2015).

The role of cytokines in the stress response and development of anxiety is not clear. Immune signaling molecules appear to be involved in stress conditions, including chronic mild stress (You et al., 2011); repeated social defeat (Wohleb et al., 2011); stress-induced ischemia (Caso et al., 2006); and HPA regulation (Turnbull and Rivier, 1999). Acute stress alone can induce peripheral cytokine production (Qing et al., 2020; Serrats et al., 2017), and cytokines are linked to the stress-induced development of anhedonia (Koo and Duman, 2008) and anxiety (Iwata et al., 2016). Tumor Necrosis Factor α (TNF) in particular may play a causal role in the development of anxiety-like behaviors, as blocking TNF signaling reduced the expression of anxiety-like behaviors induced by peripheral inflammation (Camara et al., 2015), chronic pain (Chen et al., 2013), obesity (Fourrier et al., 2019), or experimental autoimmune encephalomyelitis (Haji et al., 2012). Moreover, elevating central TNF levels by direct intracerebroventricular injection increases anxiety-like behaviors in mice (Haji et al., 2012). This suggests that it is possible that stress-induced anxiety is mediated by TNF.

A single bout of stress is reported to vary in its ability to stimulate TNF expression but generally observed to trigger an elevation (Iwata et al., 2016; Madrigal et al., 2002; Nguyen et al., 1998; O’Connor et al., 2003; Ohgidani et al., 2016; Vecchiarelli et al., 2016), similar to what is seen for chronic stress (Goshen et al., 2008; You et al., 2011). In particular, a combination of restraint and water immersion will increase TNF levels in the hippocampus due to microglia activation (Ohgidani et al., 2016). The elevation of TNF in the hippocampus is intriguing as there is strong evidence to implicate the ventral hippocampus in the regulation of anxiety-like behaviour. Unlike the dorsal hippocampus, a key structure for spatial memory, the ventral hippocampus modulates emotional states (Fanselow and Dong, 2010; Strange et al., 2014). This is particularly true for anxiety, as lesioning the ventral hippocampus is anxiolytic (Bannerman et al., 2002; Kjelstrup et al., 2002). Anxiety-responsive cells are found in the ventral hippocampus, and optogenetic activation of the terminals of these cells in the lateral hypothalamic area increased avoidance and anxiety-like behaviour (Jimenez et al., 2018). Critically, a variety of optogenetic and pharmacologic manipulation of the ventral hippocampus (and its inputs and outputs) directly alter anxiety-like behaviours (Felix-Ortiz et al., 2013; Jimenez et al., 2018; Kheirbek et al., 2013; Kjaerby et al., 2016; Padilla-Coreano et al., 2016; Parfitt et al., 2017; Wu and Hen, 2014). The ventral hippocampus has also been shown to be involved in descending regulation of the HPA in response to acute stress (Radley and Sawchenko, 2011).

However, whether the increase in hippocampal TNF is instructive to stress-related synaptic modulation and associated anxiety-like behaviors remains unexplored. In light of this, we investigated the mechanistic role for TNF, an inflammatory signaling molecule shown to play a role in the homeostasis of synaptic transmission (Stellwagen and Malenka, 2006), in the stress response. Here we demonstrate that acute stress induces TNF production within the ventral hippocampus. Further, sustained hippocampal TNF signaling is required to maintain the synaptic potentiation and the increase in anxiety-like behavior induced by stress.

## Results

We used an acute stressor to study the role of TNF on subsequent stress induced synaptic and behavioral changes (Fig 1A). Exposure of adult male mice to 20 min of forced swimming (at 23-25°C) was sufficient to increase anxiety-like behaviour 24h later as measured by both the light-dark box and open field test (Fig 1B). This effect was not stressor dependent, as 1.5hrs of restraint stress caused a similar increase in anxiety-like behaviour, measured 24h later, using the light-dark box (Fig 1C). The canonical stress pathway is necessary for this stress-induced increase in anxiety-like behaviour: inhibiting Cort production, the principal terminal output of the HPA axis, via metyrapone administration (IP; 50mg/kg; 1 h pre-stress) blocked the behavioral changes seen in the WT animals (Fig 1D).

**Figure 1:**
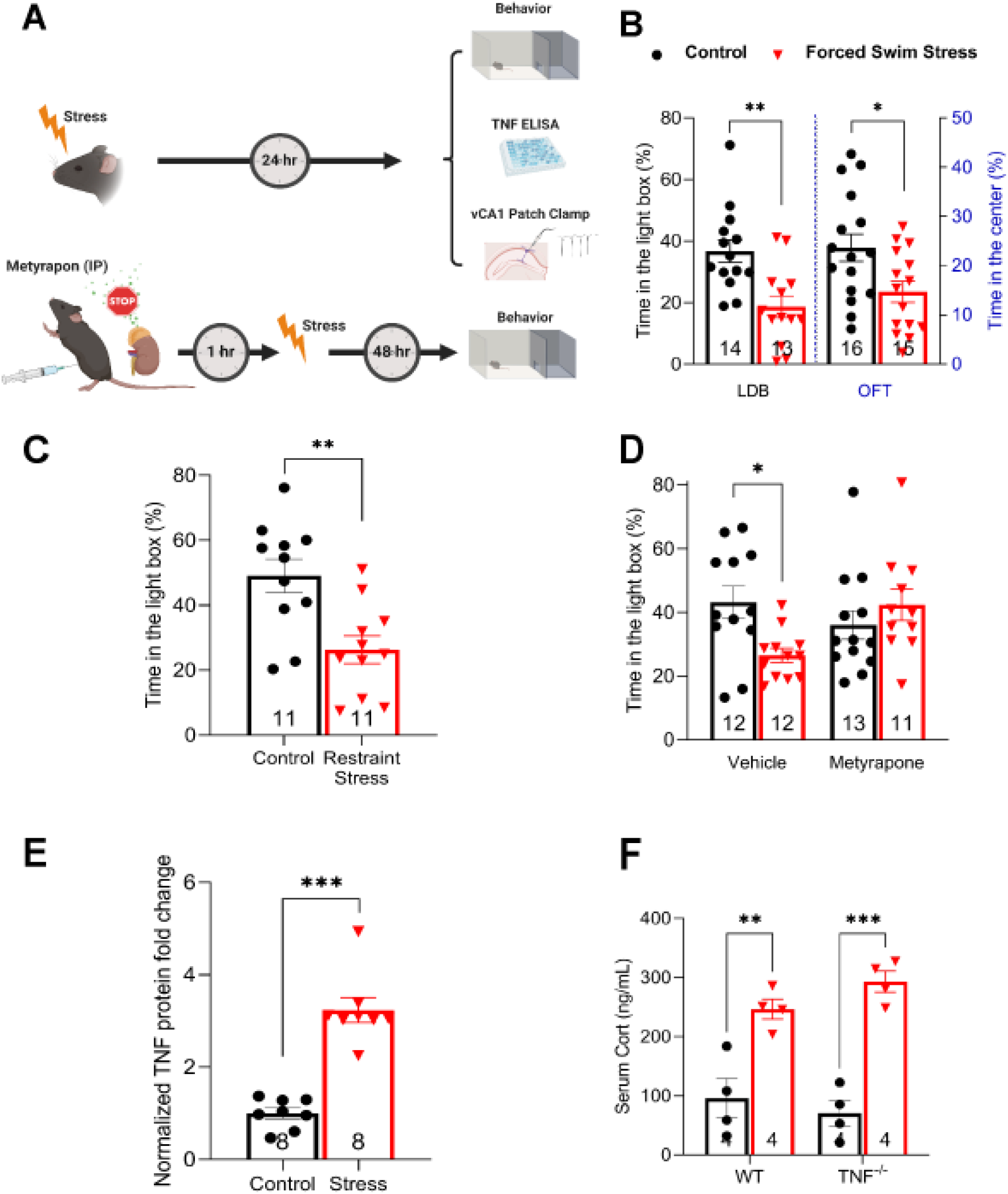
Molecular and behavioral characterization of acutely stressed mice. A. A general schematic of the conducted experiments. For most experiments, mice were subjected to a single bout of acute stress and tested 24h later. Some mice were given metyrapone 1h prior to stress and then tested 48h later. B. Animals subjected to forced swim stress display anxiety like behavior, as measured in the LDB (two-tailed student t-test P = 0.0015) and the OFT (two-tailed student t-test P = 0.0171). C. The induction of anxiety-like behavior in males is also observed in response to RS paradigm (two-tailed student t-test, P = 0.0028). D. Blocking CORT synthesis through administering metyrapone (IP; 40 mg/kg; 1 h pre-stress) inhibits stress-induced anxiety-like behavior (significant interaction of stress x drug; two-way ANOVA F (1, 44) = 7.051, P = 0.0110; Tukey’s post hoc analysis of control versus stress within vehicle P = 0.0444, within metyrapone P = 0.7281). E. Acute swim stress induces significant induction of TNF protein in the vHPC measured 24h post-stress (P<0.0001). F. Swim stress induces serum CORT levels independent of TNF signaling (ANOVA F (1, 12) = 63.79, P<0.0001; Tukey’s post hoc analysis of control versus stress within WT animals P = 0.0031, within TNF^-/-^ animals P = 0.0001) *P < 0.05, **P < 0.01, ***P < 0.001. Data are presented as mean± SEM. TNF, tumor necrosis factor; CORT, corticosterone; LDB, light-dark box. OFT; open field test. Sample sizes are indicated on the figures.

Several studies have reported that acute stress induces TNF expression in the forebrain for up to seven hours (Iwata et al., 2016; Madrigal et al., 2002; Ohgidani et al., 2016; Vecchiarelli et al., 2016), but the role for this persistent elevation remains unclear. In response to forced swim stress, we observed a robust elevation in TNF protein in the ventral hippocampus at 24h post-stress (Fig 1E).

To investigate the role of elevated TNF in the initiation or expression of the stress response, we began by comparing circulating Cort levels between wildtype (WT) and TNF deficient (*TNF^-/-^*) mice, both under basal conditions as well as shortly after exposure to acute swim stress. We found no significant differences under either condition (Fig 1F), demonstrating that TNF is not involved in regulating circulating Cort levels basally nor its release following stress. Stress is also known to be a potent activator of immediate early gene expression in the hippocampus (Cullinan et al., 1995). Synaptic output from ventral hippocampus is strongly linked to the expression of anxiety-related behaviours (Jimenez et al., 2018; Padilla-Coreano et al., 2016) and this region is involved in descending regulation of the HPA in response to acute stress (Radley and Sawchenko, 2011). Therefore, we tested whether *TNF^-/-^* animals have a normal c-Fos response to stress in the ventral hippocampus. We observed that the stress-induced increase in c-Fos expression was comparable between *TNF^-/-^* and WT animals (Supplemental Fig 1). As a whole, these data suggest Cort regulation is independent of TNF, and thus TNF would more likely be a downstream mediator of the stress response.

To test the role of TNF in the expression of stress-induced behavioral change, we next examined the difference in stress-induced changes in anxiety-like behaviour between WT and *TNF^-/-^mice.* Using the light-dark box, *TNF^-/-^* mice spent an equivalent amount of time in the light compartment as their WT counterparts under control conditions but importantly did not show any decrease in this measure following stress (Fig 2A). Mice lacking TNF also did not have the normal anxiety-induced increase in latency to enter the light compartment and decrease in transitions between the compartments (Supplemental Fig 2). Similarly, mice lacking the principal receptor for TNF (*TNFR1^-/-^*) also had normal baseline behaviour but no stress-induced elevation in anxiety-like behaviour (Supplemental Fig 3).

**Figure 2:**
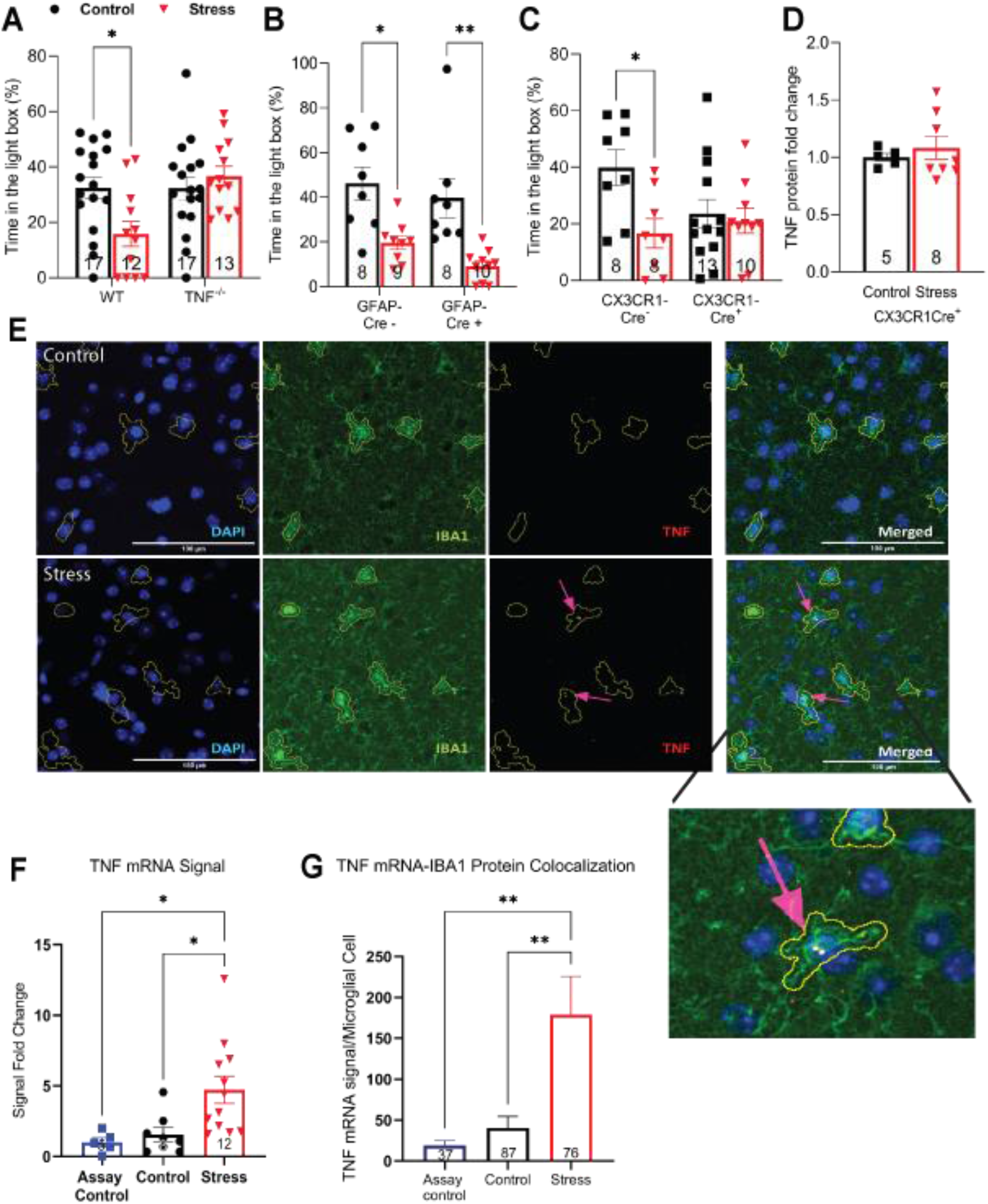
Microglial TNF signaling is important for the induction of stress-induced anxiety-like behavior. **A.** Mice that lack global TNF production do not show induction in anxiety-like behavior. (two-way ANOVA genotype x stress interaction F(1, 55) =6.536, P = 0.0134; Tukey’s *post hoc* analysis of control versus stress within WT P = 0.0324, TNF^-/-^ P = 0.8689). **B**. Stressed CX3CR1-Cre positive animals lack stress-induced increase in anxiety-like behavior compared to their unstressed counterparts. CX3CR1-Cre negative animals show a comparable phenotype to WT (24 h; two-way ANOVA main effect of stress F (1, 35), P = 0.0227; main effect of genotype F (1, 35) = 1.222, P = 0.2765; interaction of stress x genotype F (1, 35) = 3.679, p = 0.0633). Tukey’s *post hoc* reveals that for Cre negative, control versus stress P = 0.0384 while for Cre positive, control versus stress P = 0.9836). **C.** Stress induces anxiety-like behavior in GFAP-Cre positive animals compared to their unstressed GFAP-Cre positive counterparts, similar to the phenotype in WT animals (24 h; two-way ANOVA main effect of stress F(1, 31) = 25.53, p <0.0001; Tukey’s *post hoc* analysis: Cre negative control versus stress P = 0.0127, Cre positive control versus stress P = 0.0028). **D.** Microglial-TNF-lacking animals lack stress-induction of TNF in the vHPC compared to their controls (two-tailed student t-test, P = 0.5355). **E.** Sample images demonstrating the results of RNAscope *in situ* hybridization combined with IBA1 immunohistochemistry staining in the vCA1-SR, using a bacterial probe and TNF mRNA probes in unstressed controls and stressed animals. The results are quantified in F and G. **F.** Stress induces TNF mRNA in the SR of the vHPC (4 h post-stress; one-way ANOVA, main effect F(2,22), P = 0.0096; Tukey’s *post hoc* analysis of control versus stress P = 0.0288, stress versus nonspecific signal P = 0.0271, control versus nonspecific signal P = 0.9182, Sample size n (number of animals N) for assay control = 5 (3), for unstressed controls = 8 (4), for stressed animals = 12 (6)). **G.** Quantification of the overlap in *in situ* hybridization signal and microglial cells in all three conditions. (4 h post-stress; one-way ANOVA main effect F(2, 197) = 7.144, P = 0.001; Tukey’s *post-hoc* analysis of control versus stress P = 0.0029, control versus nonspecific signal P = 0.9108, stress versus nonspecific signal P = 0.0082, number of detected microglia n’ (number of sections n – number of animals N) for assay control = 37 (3-2), for unstressed controls = 87 (6-3), for stressed animals = 76 (5-3)). *P < 0.05, **P < 0.01, ***P < 0.001. Data are presented as mean± SEM. TNF, tumor necrosis factor; vHPC, ventral hippocampus. Sample sizes are indicated on the figures.

The stress-induced elevation in TNF appears to be microglial in origin. Mice with a specific deletion of TNF from astrocytes (using GFAP-Cre combined with floxed TNF alleles) have an equivalent increase in anxiety-like behaviour following stress as their Cre negative littermates (Fig 2B). However, conditional deletion of TNF from microglia using CX3CR1-Cre prevented the increase in anxiety-like behavior in stressed CX3CR1-Cre positive mice relative to their unstressed CX3CR1-Cre expressing littermates, although there was a baseline behavioural shift in these animals (Fig 2C). These data were corroborated by the lack of stress-induced elevation of hippocampal TNF in animals lacking microglial TNF (Fig 2D). Consistent with an increase in microglia activation, we also observed a stress-induced increase in IBA1 immunostaining in the ventral hippocampus and a shift to a less polarized microglial morphology (Supplemental Fig 4). We further used RNAscope to label TNF mRNA in sections from the same animals. Analysis of the stratum radiatum of the ventral hippocampus revealed a stress-induced increase in TNF mRNA labeling (Fig 2E-F) as well as an increase in TNF mRNA associated with microglia (Fig 2G). This is consistent with the stress-induced activation of microglia observed previously by others (Calcia et al., 2016; Serrats et al., 2017).

Acute stressors, including forced swim and restraint, have been shown to modify basal glutamatergic transmission in a Cort-dependent manner at several forebrain synapses (Campioni et al., 2009; Yuen et al., 2009; Yuen et al., 2011). We assessed glutamatergic transmission at Schaffer Collateral (SC) to CA1 synapses in the ventral hippocampus. Acute swim stress resulted in an increase in the AMPA/NMDA ratio at the SC synapses on CA1 pyramidal cells, relative to control mice left in their home cages. This potentiation was evident within 2h and was sustained for at least 24h (Fig 3A and Supplemental Fig 5A). Acute stress has been demonstrated to induce plasticity of AMPA receptor (AMPAR) currents, NMDA receptor (NMDAR) currents, or both, depending on brain structure examined (Campioni et al., 2009; Kuzmiski et al., 2010; Yuen et al., 2009). Given this, we pharmacologically isolated AMPAR or NMDAR currents and assessed stimulus input-current output (I-O) relationships 24 h following stress. We found that AMPAR-mediated transmission was significantly potentiated (Fig 3B) while there was no significant change in NMDAR-mediated transmission (Fig 3C). Some studies have demonstrated a pre-synaptic locus of expression of enhanced glutamatergic transmission shortly following acute stress (Musazzi et al., 2010; Treccani et al., 2014), but we observed no changes in pre-synaptic glutamate release, as measured by the coefficient of variability (CV) and paired pulse ratio (PPR) (Supplemental Fig 5B-C). Since acute stress or glucocorticoid administration have also been shown to alter expression of some intrinsic voltage-gated currents and ion channel subunits (Gray et al., 2014; Joels et al., 2003; Morsink et al., 2007), we also compared basic cellular electrophysiological properties of CA1 pyramidal cells - resting membrane potential, membrane resistance, and membrane capacitance - between stress and control groups and found them unchanged (Supplementary Fig 5D-F). This suggests that the observed potentiation of glutamatergic transmission in ventral hippocampal SC to CA1 synapses is principally due to post-synaptic AMPAR trafficking, as has been shown for prefrontal cortex pyramidal cells (Yuen et al., 2011).

**Figure 3.**
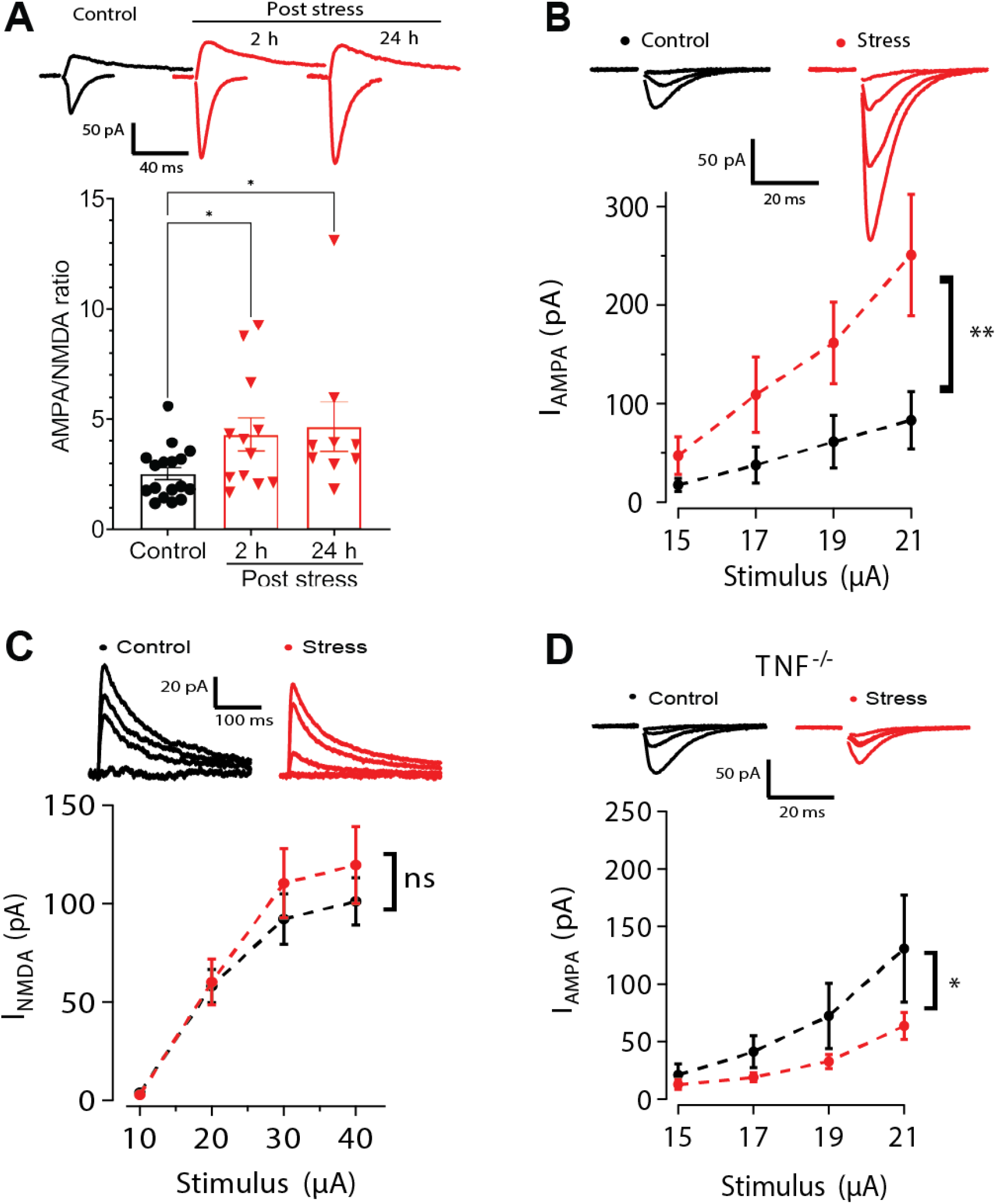
AMPAR-mediated currents at the Schaffer collateral synapses in vCA1 are potentiated in WT animals and not in *TNF^-/-^* mice. A. Acute swim stress increases the AMPA/NMDA current ratio at Schaffer collateral synapses in the vCA1 at 2 and 24 h post-stress, as seen in sample traces and group data (Kruskal Wallis H(2) = 8.068, P = 0.0177; Control: n = 17, N = 11; 2 h post-stress: n = 13, N = 8; 24 h post-stress: n = 9, N = 5). B. Stress potentiates the AMPA receptor (AMPAR) input-output relationship in the ventral Schaffer collaterals (24 h post-stress; two-way ANOVA of the main effect of stress F (1,40) = 11.439, P = 0.0016; control: n = 5, N = 4, stress: n = 7, N = 3) C. There is no significant stress-induced potentiation of the NMDA receptor (NMDAR) mediated input-output curve in the ventral Schaffer collaterals (24 h post-stress; two-way ANOVA of the main effect of stress F (1,52) = 1.0999, P = 0.2991; control: n = 7, N = 4, stress: n = 8, N = 4). D. TNF lacking animals (TNF^-/-^) do not exhibit potentiation in the AMPAR currents in the ventral hippocampus in response to stress-exposure (24 h post-stress; two-way ANOVA of the main effect of stress F (1, 44) = 6.315, P = 0.0157; control: n = 6, N = 3, stress: n = 7, N = 4). *P < 0.05, **P < 0.01, ***P < 0.001. Data are presented as mean± SEM. Sample sizes are indicated on the bar figure or in the caption.

We then tested the TNF dependence of this increase in AMPAR-mediated currents. We first determined that basal AMPAR-mediated transmission in the ventral hippocampus was not significantly different between WT and *TNF^-/-^* animals (Supplemental Fig 6A). However, *TNF^-/-^*animals did not show potentiation of AMPAR currents after being exposed to acute stress *in vivo;* instead AMPAR currents tended to be depressed following stress (Fig 3D). Stress-induced Cort release has generally been assumed to directly drive associated *in vivo* synaptic changes, in part because *ex vivo* (Karst and Joels, 2005) or *in vitro* (Groc et al., 2008; Yuen et al., 2011) application of Cort can potentiate synapses. We therefore tested the *ex-vivo* Cort response of WT and *TNF^-/-^* hippocampal slices. In marked contrast to the *in vivo* results, TNF was dispensable for the *ex vivo* Cort-induced potentiation of AMPAR current at CA1 synapses in the ventral hippocampus (Supplemental Fig 7). This suggests that *ex-vivo* application of Cort does not replicate the *in vivo* mechanism, where TNF signaling is required.

To address potential off-target effects stemming from our use of germline *TNF^-/-^* animals, we turned to a pharmacological strategy using the brain-permeant reagent XPro1595, a dominant negative version of TNF (DN-TNF) that antagonizes TNF signaling in a temporally constrained manner (Steed et al., 2003). Pre-administration of DN-TNF (IP; 20 mg/kg) prior to exposing WT mice to stress (Fig 4A) completely blocked the subsequent stress-induced potentiation of AMPAR synaptic currents (Fig 4B-C). This was specific to the potentiated synapses since DN-TNF administration under control conditions did not alter basal synaptic transmission (Supplemental Fig 6B). These data indicate that TNF is required during the stress response for the subsequent synaptic changes. As *in vivo* TNF-mediated synaptic changes are not typically long-lasting (Lewitus et al., 2016; Lewitus et al., 2014), this suggests that persistent TNF signalling might be necessary to maintain the potentiation. To address this, we first tested whether the stress-induced synaptic plasticity could be reversed following the induction and expression of the potentiation, which occurs within 2h (see Fig 3A). Indeed, we found that administering DN-TNF to animals 5-24h after swim stress (Fig 4D) was sufficient to reverse the stress-induced synaptic potentiation at 48h post-stress (Fig 4E-F).

**Figure 4:**
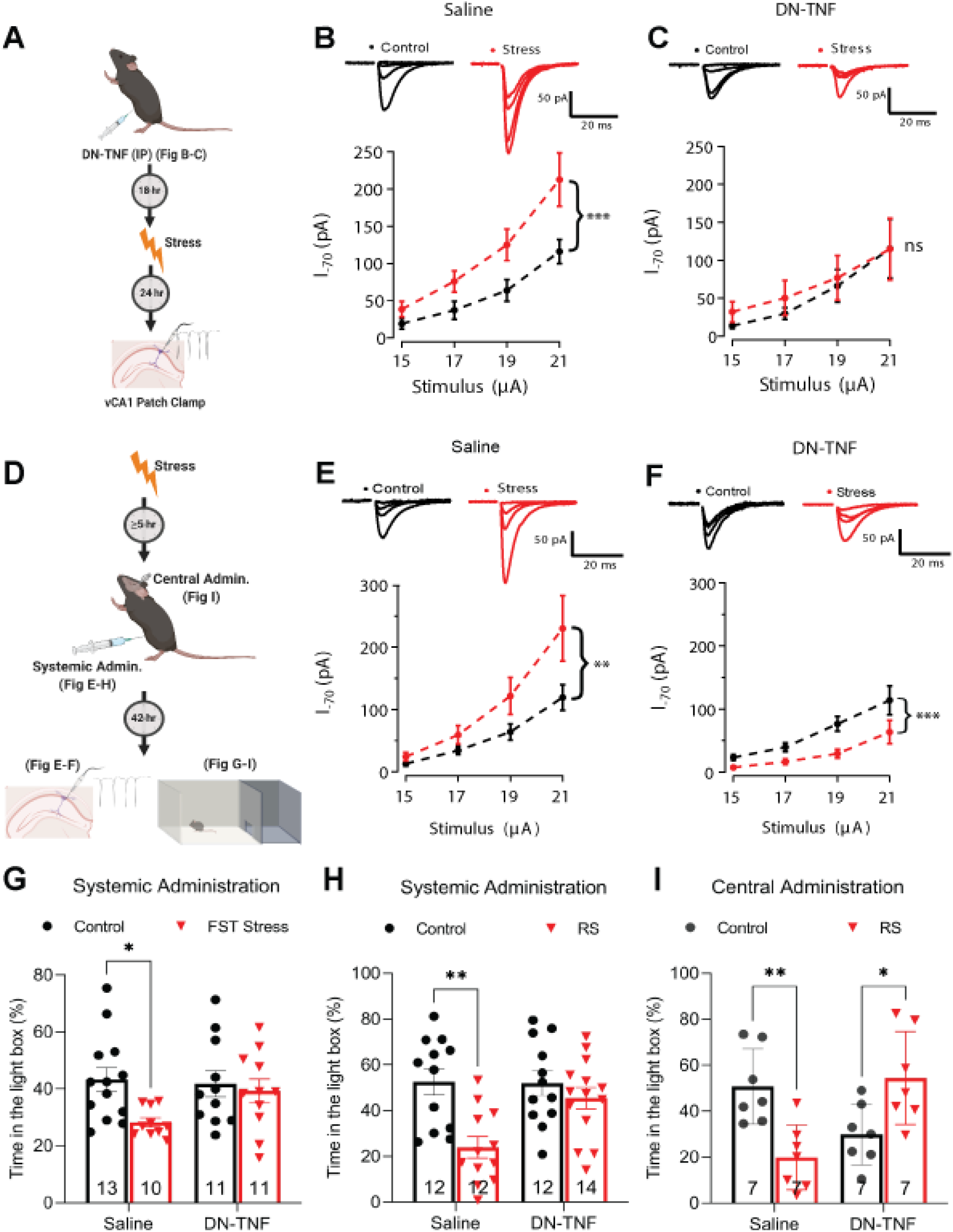
TNF antagonism post-stress reverses stress-induced plasticity and behavior. A. A schematic of the experiments performed in B and C. B. Systemic (IP) administration of saline prior to stress does not interfere with stress-induced AMPAR current potentiation at vCA1-SC synapses (two-way ANOVA of the main effect of stress F (1, 58) = 14.5093, P = 0.0003; control: n = 7, N = 5, stress: n = 10, N = 6). C. Blocking TNF signaling through systemic (IP) administration of dominant negative TNF (DNTNF; 20 mg/kg) prior to stress blocks the observed potentiation (two-way ANOVA of the main effect of stress F (1, 44) = 0.4727, P = 0.4954; control: n = 7, N = 3, stress: n = 6, N = 4). D. A schematic of the experiments performed in E-I. E. Administering saline IP at 5h post-stress does not block AMPAR mediated, vCA1-SC synaptic potentiation observed at 48h post-stress (two-way ANOVA of the main effect of stress F (1, 84) = 6.5693, P = 0.0122; control: n = 11, N = 7, stress: n = 12, N = 8). F. Systemic (IP) administration of DN-TNF 5h after stress blocks the expression of stress-induced synaptic potentiation (two-way ANOVA of the main effect of stress F(1, 64) = 14.78, P = 0.0003; control: n = 10, N = 5, stress: n = 8, N = 5). G. Blocking TNF signaling via administering DN-TNF (IP; 20 mg/kg; 5h post-stress) prevents anxiety-like behavior 48h post-swim stress (two-way ANOVA of the main effect of stress F (1, 41) = 4.825, p = 0.0338; main effect of drug treatment F (1, 41) = 1.428, p = 0.2390; interaction of stress x drug treatment F (1, 41) = 2.470, p = 0.1237; Tukey’s post hoc analyses of control saline versus FST saline p = 0.0494, control DN-TNF versus FST DN-TNF p = 0.9714). H. TNF is similarly necessary for stress-induction of anxiety-like behavior following restraint stress (two-way ANOVA of the interaction of stress x drug treatment F (1, 46) = 4.692, p =0.0355; Tukey’s post hoc analyses of control saline versus RS saline p = 0.0017, control DN-TNF versus RS DN-TNF p = 0.7908). I. Blocking TNF in the vHPC through local administration of DN-TNF (IC; 4.8 mg/kg; 5h post-RS) prevents the development of anxiety-like behavior 48h in response to stress (two-way ANOVA of the interaction of stress x drug treatment F (1, 24) = 20.58, p =0.0001; Tukey’s *post hoc* analyses of control saline versus RS saline p = 0.0078, of control DN-TNF versus RS DN-TNF p = 0.0419). *P < 0.05, **P < 0.01, ***P < 0.001. Data are presented as mean± SEM. Sample sizes are indicated on the bar figure or in the caption.

We then proceeded to test the behavioral consequences of this protocol. As with the synaptic potentiation, administration of DN-TNF 5h after swim stress completely abrogated the anxiety-like phenotype seen in the saline treated cohort (Fig 4G). These results were replicable using restraint stress, as administering DN-TNF 5h after this stressor also blocked the stress-induced increase in anxiety-like behavior (Fig 4H). This was dependent on TNF signaling within the ventral hippocampus, as local injection of DN-TNF bilaterally into the ventral hippocampus was sufficient to block the expression of stress-induced anxiety-like behavior relative to littermates injected with saline (Fig 4I). Neither stress, drug, nor mode of administration impacted the locomotor behaviour of the animals (Supplemental Fig 8). We therefore conclude that acute stress induces on-going TNF signalling within the ventral hippocampus that is required to maintain the synaptic and behavioral outcomes of the stressor, in particular anxiety-like behaviours.

## Discussion

This TNF-mediated potentiation correlates with the stress-induced increase in anxiety like behavior, and blocking TNF-signaling even hours after the stressor reverses both the synaptic potentiation and anxiety like behavior.

Our results here reveal a novel property of the *in vivo* stress response, dissecting the response into at least two distinct phases: an induction phase and a maintenance phase. Importantly, we did not observe any gross abnormalities in the neuroendocrine axis of TNF deficient animals, nor in the stress-induced expression of the immediate early gene *c-Fos*, again highlighting that the deficit these animals exhibit with respect to the stress response appears limited to a lack of expression of a specific form of synaptic plasticity.

The induction phase is likely Cort dependent, as blocking Cort production with metyrapone prevented the stress-induced anxiety. Previous results have also implicated Cort in the synaptic potentiation (Campioni et al., 2009; McEwen et al., 2015; Popoli et al., 2011; Yuen et al., 2009). However, the majority of work suggesting that Cort directly drives the potentiation comes from in vitro application of Cort (Groc et al., 2008; Karst and Joels, 2005; Yuen et al., 2011). While we can replicate the substantial potentiation of hippocampal synapses by Cort *in vitro* (Supplemental Figure 7), other work suggests that this potentiation is not sustained beyond a few hours (Karst and Joels, 2005). Further, slices from *TNF^-/-^* mice display an equal Cort-driven synaptic potentiation *in vitro*, but no sustained stress-induced hippocampal potentiation *in vivo.* Thus we would argue that Cort is necessary to induce the stress response but the sustained synaptic potentiation *in vivo* is maintained by on-going TNF signaling.

Consistent with our findings, TNF can directly increase AMPAR content at CA1 hippocampal synapses (Beattie et al., 2002) and increased output from the ventral hippocampus can drive anxiety-like behavior (Jimenez et al., 2018). Thus TNF is likely to drive the observed synaptic potentiation on CA1 pyramidal neurons, which may well be accompanied by decreased inhibition (Stellwagen et al., 2005), and would result in increased activity of these cells. That blocking TNF signaling just within the ventral hippocampus is sufficient to prevent the stress-induced anxiety would argue that these synaptic changes in the ventral hippocampus are causative for the stress-induced increase in anxiety-like behavior.

Our data here also supports the idea that microglia are the source of the stress-induced elevation in TNF. Consistent with previous reports (Calcia et al., 2016; Ohgidani et al., 2016; Serrats et al., 2017), we observe a stress-induced increase in markers of microglia activation, including increase Iba1 expression and altered microglia morphology. Moreover, we observed an increase in TNF mRNA associated with microglia. The conditional deletion of TNF from microglia prevented the stress-induced increase in hippocampal TNF, and also prevented a stress-induced increase in anxiety-like behavior. However, we did observe a baseline shift in anxiety-like behavior in these mice, which was not seen in either the full TNF knockout or the astrocyte-specific knockout. This complicates the interpretation, and we have observed several changes in baseline behavior in the microglial-conditional mice (including increased aggression and fighting) that could drive other changes in baseline behavior. But despite the baseline shift, there should be adequate potential to express additional anxiety-like behavior as the mice are still spending over 20% of their time in the light compartment, and thus we don’t believe that any floor effect is obscuring a stress-induced change in behavior. Importantly, when combined with the other data on stress-induced microglial activation and lack on TNF response to stress in the conditional mutant, we believe that a microglia source of stress-induced TNF is the most parsimonious hypothesis. Whether the microglial activation is due to Cort signaling is unclear at this point. Our data suggests that Cort signaling is critical for the stress-induced behavioral change, but the direct activation will need to be tested. Microglia express both MR and GR (Brocca et al., 2019; Chantong et al., 2012), and stress can act through GR to increase TNF levels (Frank et al., 2012; Wang et al., 2017). A glial target of Cort signaling would be consistent with the finding that conditional deletion of GR from forebrain excitatory neurons did not alter a stress-induced increase in anxiety-like behavior (Furay et al., 2008).

Other cytokines may also be released by activated microglia, and there is evidence that IL1 contributes to stress induced anxiety-like behavior (McKim et al., 2018), although the observed increase in IL1 was more transient than the increase in TNF. This could suggest that multiple cytokines are involved with the induction of the stress response but perhaps TNF has a unique role in maintaining the anxiety.

One interesting observation is the significant decrease in synaptic strength post-stress observed in *TNF^-/-^* and DN-TNF treated animals, which suggests that the synaptic strength is actually decreasing in response to stress. This might be indication of a countervailing resilience-type signal being unmasked in the absence of TNF signaling, though the nature of such a signal would need to be determined.

Critically, the functional consequences of this stress-induced potentiation can be reversed well after its onset by blocking TNF signaling. The synaptic potentiation is established within 2 h of the stressor, yet blocking TNF signaling 5 h afterwards is sufficient to reverse both the synaptic plasticity and the behavioral change. Thus hippocampal TNF signaling is necessary for the maintenance of these changes. This may have therapeutic potential for cases in which the effects of stress are particularly detrimental as in post-traumatic stress disorder, where TNF-based therapeutics may hold potential to treat the chronic anxiety of such conditions.

## Materials and Methods

All experiments and procedures involving animal use were conducted in accordance with the guidelines of the Canadian Council on Animal Care and approved by the local Animal Care Committee.

### Animals

All experiments were done using adult C57BL/6 male mice, 8-16 week old for behavioral experiments and 8-12 week old for electrophysiology experiments. TNF homozygous knockout (*TNF^-/-^*) mice, obtained from Jackson Labs, were bred in house, and compared with wild-type (WT) mice of the same background strain (C57BL/6J) also bred in house. The in-house WT and *TNF^-/-^* lines were periodically backcrossed. In some cases, WT C57BL/6 mice were obtained from Charles River, in which case they were housed in the animal facility for at least one week prior to experimentation*. TNFR1^-/-^* mice (Peschon et al., 1998) were obtained from Jackson Labs and bred homozygously. Floxed TNF mice were obtained from S. Nedospasov (Grivennikov et al., 2005) and crossed with GFAP-Cre mice (Bajenaru et al., 2002) from NCI Mouse Repository (RRID: IMSR_NCIMR: 01XN3) or with Tg(Cx3cr1-cre)MW126Gsat mice (Yona et al., 2013) generated by N. Heintz (The Rockefeller University, GENSAT) and purchased from MMRRC (UC Davis; RRID: MMRRC_036395-UCD). GFAP or Cx3Cr1-Cre expressing mice were compared with GFAP or CX3CR1-Cre non-expressing littermates. Floxed TNF and GFAP-Cre mice were on a C57BL/6 background; Cx3Cr1-Cre mice were a mix of FVB/B6/129/Swiss/CD1 and backcrossed several generations with C57BL/6J. Animals were group housed (3-5 animals/cage, except following stress; see below) in standard controlled housing environment of 12 h light cycle, and food and water available ad libitum.

### Stress paradigms

Two stressors were used to verify the robustness of the phenotype: most animals were exposed to a forced swim of 20 min duration; the time of day in which the animals were exposed to stress was consistent throughout all experiments (10:00 AM – 1:00 PM). Animals were placed in a 4 L glass beaker, filled to the 3 L mark with room temperature (23-25 °C) tap water. At completion, excess water was gently wiped off animals and they were housed individually thereafter, with food and water available ad libitum until experimentation. Other animals were exposed to immobilization stress through restraining them in 50ml conical tubes (Sarstedt #62.547.205), perforated to allow for airflow. Tubes were then left undisturbed for 1.5 h at room temperature under regular room white fluorescent lighting conditions. Animals were again individually housed post-stress. There was no significant effect of short-term single housing on TNF protein levels in the vHPC (Supplemental Fig 9A) or on anxiety-like behavior (Supplemental Fig 9B), so single-housing does not constitute a significant stressor on its own.

### Cannula implantation and local injections

Bilateral cannulation was performed on 8-week-old mice with a recovering period of three-four weeks to allow inflammation to subside. Animals were administrated an analgesic (Carprofen; SC, 20mg/kg) 2hr before the surgery then anaesthesia was induced at 3% isoflurane and maintained at 2% throughout the surgery. Animals were injected with 1ml of warm saline at the beginning and the end of the surgery to prevent dehydration and were kept on an active heating pad to reduce anaesthesia induced hypothermia (PhysioSuite, Kent Scientific). Eye lubricant was repeatedly applied throughout the surgery to prevent eye damage. Cannulae were stereotaxically implanted into the ventral hippocampi at coordinates (relative to Bregma) of AP −3.31mm, ML ±3.47mm, DV −4.6mm; and secured with dental cement.

The skin was closed with braided silk sutures (Medtronic, KENS1173) and cannula caps were placed to prevent clogging. Animals were allowed to recover in heated cages, and administrated Carprofen (20mg/kg/day) for three days. Post-op animals were group housed for 3-4 weeks, and then exposed to 1.5hr restraint stress and single housed from then on. DN-TNF or saline was administrated 5 hr post-stress, using an injection cannula attached to a Hamilton syringe and a micropump at a rate of 0.2 μl/min to inject 0.625μl in each hemisphere. The injection cannula was then kept in place for 1-2 minutes before removal to prevent backflow. The injection apparatus was checked before and after each injection to ensure proper flow. Animals were returned to their cages and then behaviorally tested 48 hr post-stress. Injection sites were verified by histology to be in the ventral hippocampus for all animals; a representative injection site is show in Supplemental Fig 10.

### Electrophysiology

Synaptic glutamatergic currents were measured from animals that were housed under control conditions or at 2-24 hr post-stress (AMPA/NMDA ratios) or 24 h post-stress (I/O curves). A few cells for I/O curves were sampled at 2 hr post-stress, but since there was no difference in the potentiation of AMPA/NMDA ratios or AMPAR-mediated currents observed 2 h and 24 h post-stress, data from these time points were pooled. Animals were anesthetized by isoflurane inhalation, and rapidly perfused through the left cardiac ventricle with ice-cold solution (composition in mM: glycerol 222, D-glucose 11, NaHCO_3_ 26, KCl 2.5, NaH_2_PO_4_ 1.25, MgCl_2_ 3.5, MgSO_4_ 1, CaCl_2_ 0.25, Na pyruvate 1.5, Na ascorbate 0.4) pre-saturated with 95% O_2_/5% CO_2_. The brain was then removed and horizontal slices of 300-400 μm, containing the ventral hippocampus, were cut on a vibrating slicer (Leica VT1200S) in the same solution used for perfusion, followed by recovery of slices at 32 ± 2 °C for 30-45 min in solution of composition (in mM): NaCl 118, D-glucose 11, NaHCO_3_ 26, KCl 2.5, HEPES 15, NaH_2_PO_4_ 1.25, MgSO_4_ 1, CaCl_2_ 2, myo-inositol 3, Na ascorbate 0.4, saturated with 95% O_2_/5% CO_2_; slices were maintained at room temperature thereafter, and were only used for experiments after a minimum of 1 h incubation post-slicing.

Immediately prior to placing brain slices in a perfusion chamber on the stage of an upright Olympus microscope (BX51WI) equipped with IR-DIC optics, a cut was made between CA3 and CA1. Using Clampex to run Multiclamp 700 with Digidata 1440 (Molecular Devices), voltage-clamp recordings at holding potential of −70 mV (unless otherwise specified) were acquired, at 10 kHz sampling with 2 kHz low-pass filtering, from visually identified CA1 pyramidal cells using borosilicate glass pipette electrodes, pulled to produce a resistance of 2-5 MΩ when filled with internal solution of composition (in mM): CsMeSO_4_ 125, CsCl 10, HEPES 15, MgATP 4, Na_3_GTP 0.4, Na2-phosphocreatine 5, Trisphosphocreatine 5, EGTA 0.6, CaCl_2_ 0.05, pH adjusted to 7.2 with CsOH. Bath solution containing (in mM): NaCl 110, D-glucose 11, NaHCO_3_ 26, KCl 2.5, HEPES 15, NaH_2_PO_4_ 1.25, MgSO_4_ 1, MgCl_2_ 3, CaCl_2_ 4, myo-inositol 3, and saturated with 95% O_2_/5% CO_2_ was perfused at a rate of 4-8 mL / min, and maintained at 30 ± 2 °C. In all experiments, the bath solution was supplemented with 100 μM picrotoxin to block GABAAR-mediated currents; 50 μM D-AP5 was added to isolate AMPAR currents, while 15 μM NBQX was added and the holding potential adjusted to +40 mV to isolate NMDAR currents. To evoke synaptic responses, a stimulus isolator (ISO-FLEX, A.M.P.I.) was used to deliver constant current stimuli (0.1 ms pulse duration) to stratum radiatum, using either a chlorided silver wire housed in a glass pipette filled with bath solution, or, in the case of obtaining input-output relationships, matrix electrodes fabricated by FHC (MX21AES(DH1)). Care was taken in the case of obtaining input-output relationships to systematically place the stimulating electrode within ~20 μm of the cell being recorded, and control condition recordings were consistently interleaved throughout the period of experimentation. Stimuli were delivered at intervals of 5-15 s, and 6-30 sweeps were averaged for any given stimulus level (using Clampfit 10). AMPA/NMDA ratios were obtained by dividing the peak of the evoked current generated while the cell is held at −70 mV by the mean level of the current 50 ms from the onset of stimulus while the cell is held at +40 mV. Alternatively, a subset of cells was perfused with 50 μM D-AP5 while held at +40 mV, and the resultant isolated AMPAR current obtained following D-AP5 perfusion was subtracted from the compound current prior to perfusion with D-AP5 to obtain the NMDAR component of the evoked response, in which case the peak of the AMPAR current was divided by the peak of the resultant NMDAR current.

### Drug treatments

For experiments involving corticosterone treatment, corticosterone stock prepared in DMSO was diluted to 100 nM and applied to brain slices in the solution used for recovery from vibratome slicing, for 20 min at 30 ± 2 °C, with constant aeration with 95% O_2_/5% CO_2_. Slices were then washed with corticosterone-free solution, and then incubated in slice recovery solution. Voltage-clamp recordings were obtained between 2-3 h following corticosterone treatment. Controls were likewise treated, but with vehicle (0.01% v/v DMSO). Metyrapone was obtained from Tocris Bioscience #3292/50 and reconstituted to 23 mg/ml with DMSO and administrated intraperitoneally at a final doze of 50 mg/kg. TNF dominant-negative inhibitor XPro1595 (DN-TNF; Xencor, Inc.) was diluted in sterile 0.9% w/v NaCl to 2 mg/mL (intraperitoneal injection) or 5 mg/ml (direct injection into the hippocampi) and administered at a dose of 20 mg/kg. Controls were administered equivalent volumes of DMSO or sterile saline 0.9% w/v NaCl, respectively.

### Cort ELISA

Stress-induced serum corticosterone measures were obtained 15 min following completion of forced swim protocol. Animals were anesthetized by isoflurane inhalation, decapitated, and trunk blood collected in polypropylene centrifuge tubes. Samples were stored at room temperature for 1.5 h to allow for clot formation, followed by centrifugation (300 x g) to collect supernatant for storage at −80 °C until assayed. An ELISA kit from Cayman for corticosterone was used according to the suppliers’ instructions. Samples were diluted 100-fold prior to measurement.

### TNF ELISA

Bilateral ventral hippocampal tissue was collected from stressed animals (24 hours post forced swim stress) and their age matched unstressed controls. They were anesthetized by isoflurane inhalation and decapitated for a rapid removal of the brain and gross dissection of the ventral hippocampi (<1 min). Tissue was then snap-frozen in liquid nitrogen. Tissue was then immediately homogenized using a handheld homogenizer for 5 seconds on ice in 400μl of sterile PBS buffer containing a 1x protease inhibitor cocktail from Bioshop (PIC002.1). Samples were then centrifuged at 11,000 rpm for 20 minutes at 4 °C and the supernatants collected. TNF protein levels were measured according to the suppliers’ instructions of mouse TNF ELISA kit (eBioscience, Mouse TNF alpha ELISA Ready-SET-Go! Kit #88-7324). For standardization across samples, all TNF concentrations were divided by their corresponding crude-protein input concentrations as measured following the instructions of the bicinchoninic acid assay (BCA) kit from Thermo Fisher Scientific (#23227).

### c-Fos Immunohistochemistry

One hour following stress, animals were anesthetized by isoflurane inhalation, followed by transcardial perfusion with 4% w/v formaldehyde in PBS. The brain was then dissected and immersed in 4% w/v formaldehyde for 24 h at 4 °C, then transferred to 30% w/v sucrose solution. The brains were subsequently mounted in Optimal Cutting Temperature (OCT) medium to cut 30 μm slices using cryostat microtome. Slices were incubated in a blocking solution, composition: 2% v/v normal goat serum and 1% v/v affinipure donkey anti-mouse IgG (Jackson Immunoresearch) in PBS for 4 h at room temperature. Slices were subsequently incubated with primary antibodies: anti-c-Fos at 1:100 dilution (Santa Cruz Biotechnology, catalogue sc-52) and anti-NeuN at 1:200 (Millipore, catalogue MAB377) in blocking solution overnight at 4 °C. Slices were then washed twice (15 min each wash) with PBS + 0.05% Tween 20 (PBST), and once in PBS for 15 min, and incubated with secondary antibodies: Alexa568-conjugated anti-rabbit IgG at 1:300 dilution and Alexa488-conjugated anti-mouse IgG at 1:400 dilution in blocking buffer for 2 h at room temperature. Slices were then washed twice (15 min each wash) with PBS + 0.05% Tween 20 (PBST), and once in PBS for 15 min, followed by mounting in Fluoromount-G medium (Southern Biotech). Images were acquired with an Olympus Fluoview FV1000 confocal microscope using 20X objective. 25-30 optical sections at 1 μm optical sectioning were collected, and Z-projected into a single image. Multiple images of region CA1 and DG were collected for each animal, and the number of c-Fos immunoreactive cells were manually counted, and averaged for all slices to give one mean value for each animal. For clarity, the c-Fos-labeled sections used as example images shown were background subtracted and despeckled using ImageJ (NIH), followed by conversion to binary format and dilation, and then overlaid with corresponding NeuN-labeled sections.

### IBA1 Immunohistochemistry and microglial analysis

Animals were perfused 4hr post-restraint stress with 4% PFA in PBS, and whole brains post-fixed overnight (~16 hr) before being cryoprotected in 30% sucrose in PBS for 4 hr. Coronal 25 μm sections containing the ventral hippocampus were cut on a cryostat, mounted on charged slides (Fisher Scientific, catalogue 12-550-15) and dried in a desiccating chamber at room temperature for 2 hr. Sections were then washed 3x with PBS at room temperature (10 min) followed by 1 h permeabilization using 2% triton X-100 in PBS. Sections were blocked for 2 hr with 10% normal goat serum, 2% BSA and 0.3% Triton X-100, and then incubated overnight at 4°C with IBA1 polyclonal rabbit antibody (FUJIFILM Wako

Chemicals, catalogue 019-19741; 1:500) in blocking buffer. Sections were then washed as before and incubated in goat anti-rabbit secondary antibody (Thermofisher, A21244), washed 4x, and nuclei counterstained with 1:10000 of Hoechst 33342 (Thermofisher H1399) before mounting with Fluoromount-G (Thermofisher, 00-4958-02). Stacked Images were acquired using FV1000 Olympus confocal microscope using 20x objectives of the vCA1 region, with multiple images per animal. Images were then analyzed for signal intensity, cell count and morphology semi-automatically using ImageJ (NIH) and its associated FracLac package (Karperien). Average values were calculated per field, with the researcher blinded during both acquisition and analysis. The images were converted to grayscale, unsharpened, despeckled, thresholded automatically and converted to a binary mask. The “analyze particles” function was then applied with size thresholding of 50 pixels on the lower limit. The obtained ROIs were then checked manually for accuracy then were overlayed over the original image to obtain the reported outputs. Microglia morphology was evaluated using a hull and circle analysis, by fitting each microglia with an encompassing oval/circle and a convex hull. Morphology was then assessed by measuring the polarity of these shapes as a ratio of the major and minor axes and hull circularity as the hull area over the perimeter, with a loss of polarity representing an increase in activation.

### Behavior

All behaviour experiments were verified using multiple cohorts and replicated across different time intervals. Animals were tested 24-48 h following stress, and experiments were conducted between 10:00 AM and 5:00 PM (anxiety) or 4:00-7:00 PM (locomotion). Anxiety-related behavior was assessed using the Light-Dark box, which consisted of an uncovered compartment (light compartment) made of clear plexiglass, measuring 29 cm length x 20.4 cm width x 22 cm height, connected via a small box-floor level opening to a black, lid-covered compartment (dark compartment) measuring 15.5 cm length x 20.4 cm width x 22 cm height. The animal was placed in the dark compartment with the animal allowed to freely explore both compartments for a period of 10 min while video recording from directly above. The test was conducted under normal room white fluorescent lighting. The total time spent in the light compartment was scored semi-automatically using Ethovision XT 8.5 (Noldus). Anxiety-like behavior was also assessed using the Open Field Test using a 60 x 60 cm clear plexiglass box under normal room lighting conditions. The time spent in the center (out of the total testing time of 10 min) was quantified automatically using the pre-existing template in Ethovision. For assessing locomotion, proxied by the total distance travelled during a period of 20 min, animals were placed individually in 30 x 30 cm plexiglass boxes recorded from above under low intensity red light conditions. The total distance travelled during the 20 min period was then recorded automatically using Ethovision.

### Statistical analysis

All data are presented as means ± SEM. In electrophysiology data, n represents the number of cells, while N represents the number of animals; no more than three cells were recorded from any given animal, with all experimental groups being represented by at least three animals. Statistical analyses were done using Graphpad Prism 9. Data was analyzed using one or two way ANOVAs, followed by Tukey post-hoc tests, or by Student’s unpaired two-tailed t-tests. For Fig 3A, the data was not normally distributed, so we used a Kruskal-Wallis One Way ANOVA on Ranks followed by Dunn’s method of comparison with control.

## Acknowledgements

This work was supported by the Canadian Institutes for Health Research and Natural Sciences and Engineering Research Council of Canada. G.M.K was supported by a fellowship from the Canada First Research Excellence Fund, awarded to McGill University for the Healthy Brains for Healthy Lives initiative. H.F.A. was supported by doctoral awards from the Heart and Stroke Foundation of Canada, and the Research Institute of the McGill University Health Centre. We kindly thank Xencor for the donation of XPro1595. Schematic figures were created using BioRender.

---

**Supplementary 1.**
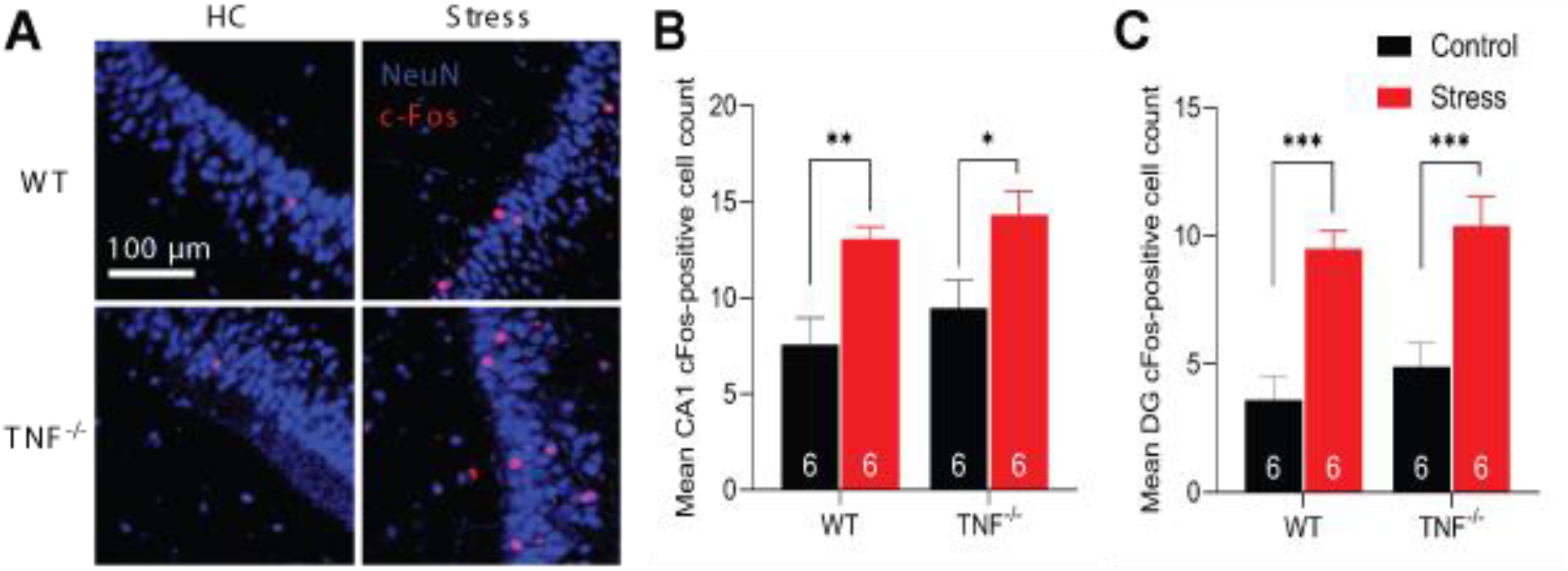
**S1. A** c-Fos labelling of the CA1 of the vHP in control and stressed animals. Labelling was significantly increased in the CA1 (**B**) and dentate gyrus (**C**) in both WT (P < 0.001) and TNF^-/-^ (P < 0.001) animals following stress. There were no statistically significant differences between genotypes under control conditions (P = 0.34), nor following stress (P = 0.51).

**Supplementary 2.**
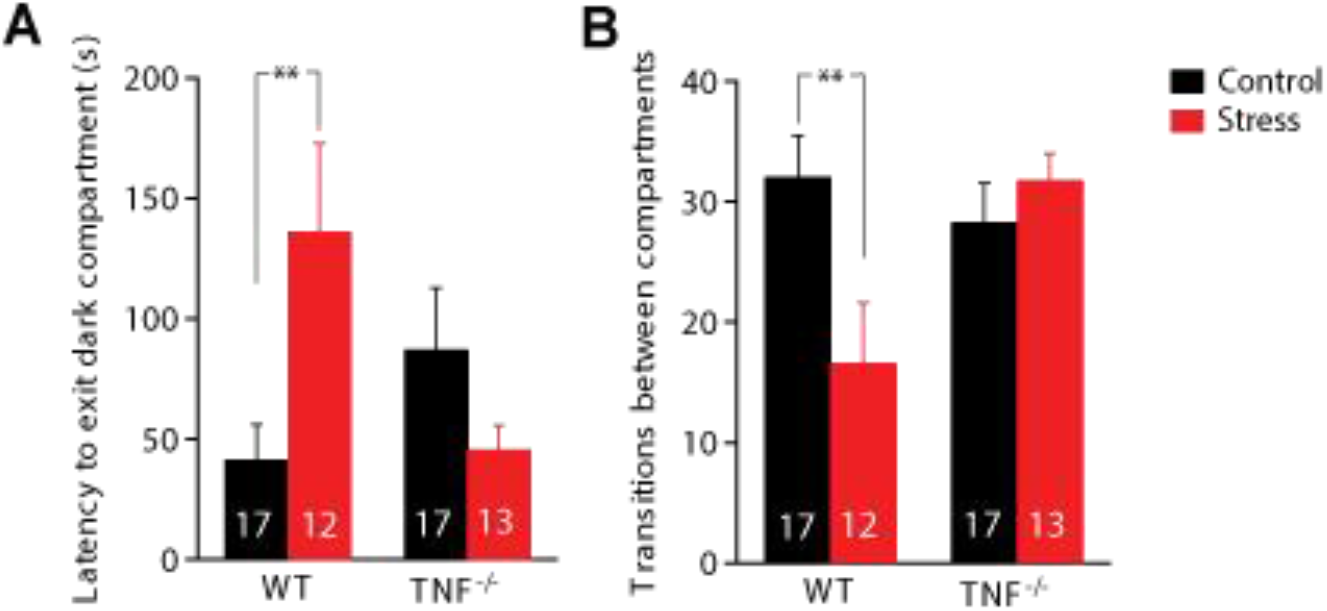
**S2. A.** Latency to first exit in the light-dark box is longer in stressed wild-type (WT) animals compared to TNF knockout (TNF^-/-^) mice (two-way ANOVA genotype x stress interaction F (1, 49) = 9.140, P = 0.0040; Tukey’s *post hoc* analysis of control versus stress within WT p = 0.0376, TNF^-/-^ p = 0.4996; Tukey’s *post hoc* WT versus TNF^-/-^ within controls p = 0.3646). **B.** Stress reduces the number of transitions WT animals make from the dark box to the light box, while it does not affect the same parameter in TNF^-/-^ mice (two-way ANOVA genotype x stress interaction F (1, 55) = 6.992, P = 0.0107; Tukey’s *post hoc* analysis of control versus stress within WT p = 0.0195, TNF^-/-^ p = 0.8990).

**Supplementary 3.**
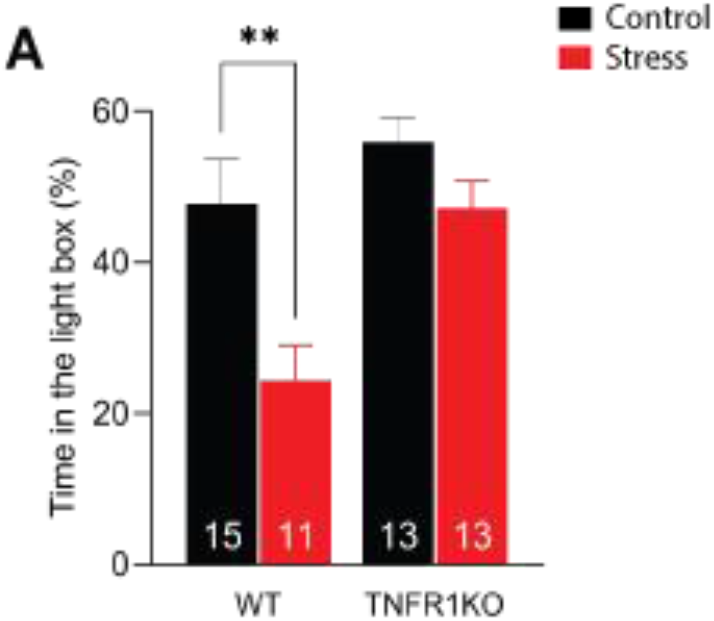
**S3.** No stress-induced anxiety-like behavior was observed in TNFR1KO animals (24 h post-stress; two-way ANOVA main effect of genotype F (1, 48) = 10.97, p = 0.0018, main effect of stress F (1, 48) = 11.73, p = 0.0013, genotype x stress interaction F (1, 48) = 2.434, p = 0.1253. Tukey’s post hoc analysis: difference within WT p = 0.0054, difference within TNFR1KO p = 0.5509). *P < 0.05, **P < 0.01, ***P < 0.001. Data are presented as mean± SEM. Sample sizes are indicated on the figures.

**Supplementary 4.**
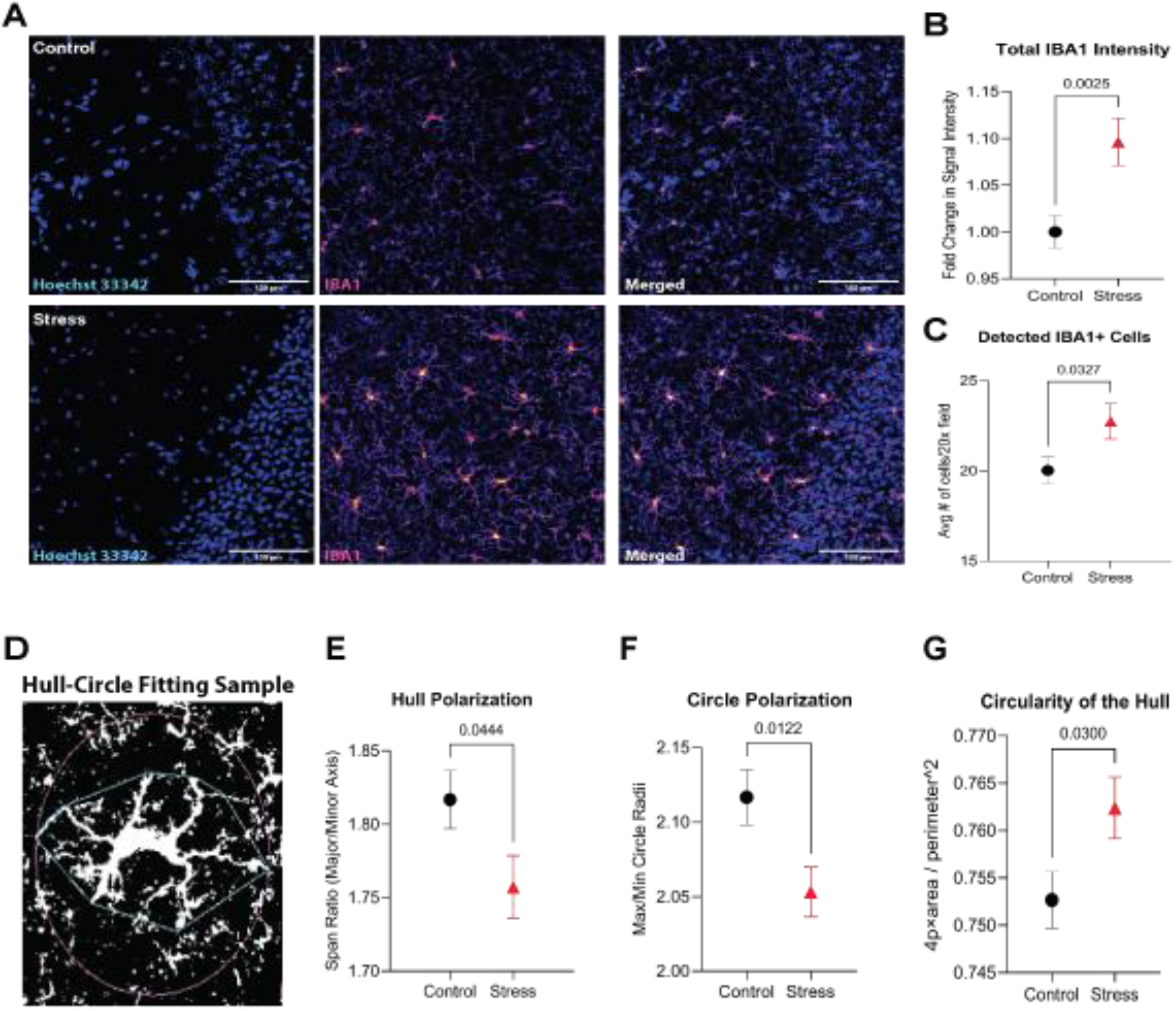
**S4. A.** Sample images demonstrating microglial activation as proxied by IBA1 signalling and cell density. **S4. B.** Quantification of total IBA1 signa in the vHPC-CA1 region of unstressed controls and acutely stressed animals (4 h post-stress, student t-test, p = 0.0025, sample size n (number of animals N) for control = 43 (4), for stress = 45 (4)). **S4. C.** Quantification of microglia in control versus stress (4 h post-stress, student t-test, p = 0.0327). **S4. D. & E.** An illustration of the pipeline for fitting microglia in a hull (inner green shape) and a circle (outer purple shape) for quantitative analysis of microglial morphology. The convex hull of a cell is the smallest convex set that encompass the cells. A convex set is a polygon where the line between any two points lies completely within the polygon. Microglia was delineated semi-automatically and the hull and circle shapes were fitted automatically using Frac-Lac package in ImageJ.(Karperien, 1999-2013.) **S4. F-H.** Quantification of three parameters of the circularity of microglia, namely the polarization of the fitted hull (p = 0.0444), the polarization of the fitted circle (p = 0.0122), and the circularity of the hull (p = 0.03, all parameters were analyzed using a student t-test, n (N) for control = 43 (4), for stress = 45 (4), see the y axes for the equations used to measure each variable). vHPC: ventral hippocampus, n: sample size, N: number of animals

**Supplementary 5.**
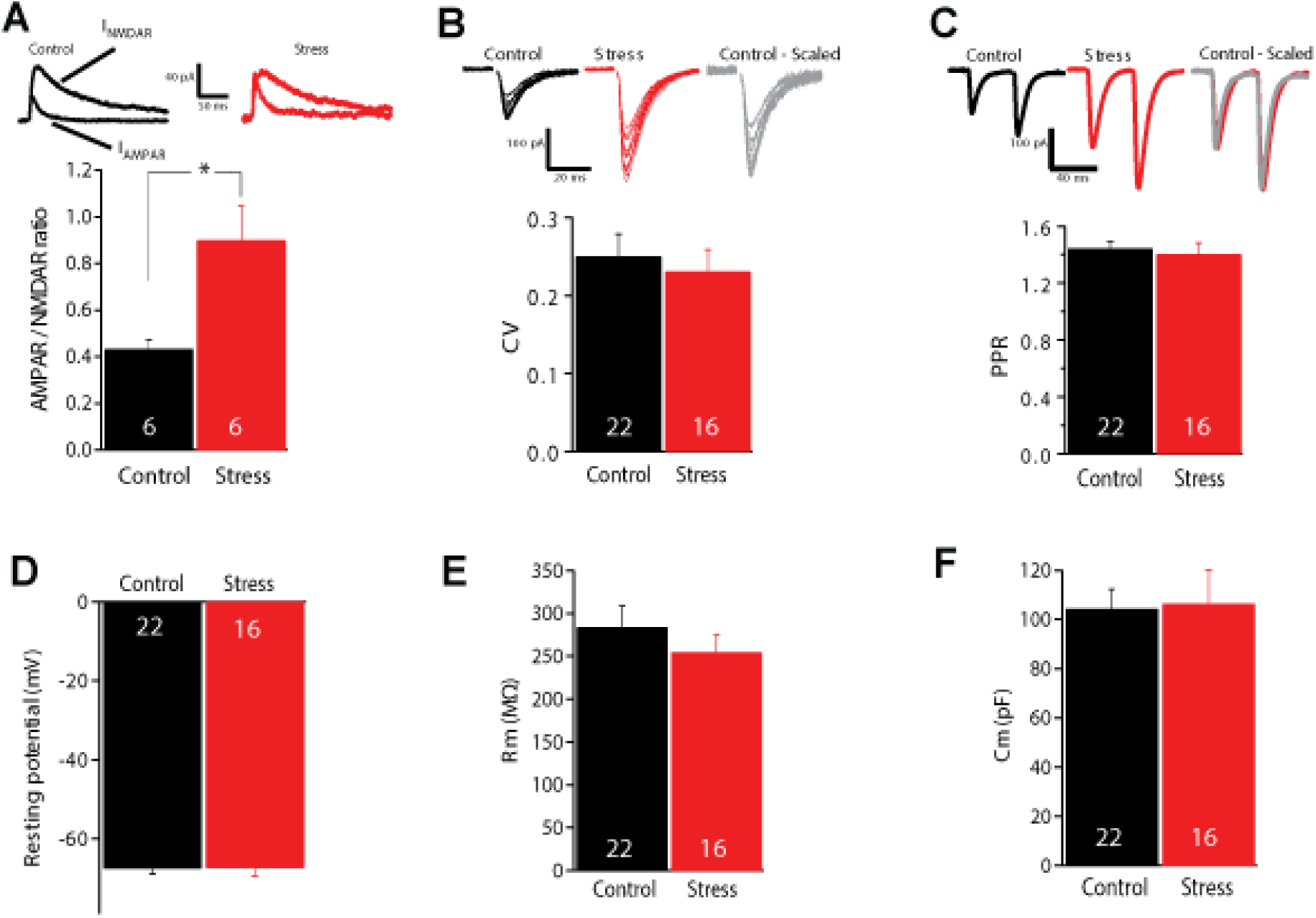
**S5. A.** Cells in the baseline and 24 h post stress groups were perfused with D-AP5 subsequent to depolarization to +40 mV to isolate the AMPAR current. Example traces illustrate the isolated AMPAR current (black trace) which was subtracted from the compound current (not illustrated) to produce the resultant (isolated) NMDAR current (red trace). The peaks of the AMPAR current and NMDAR current were used in calculation of the AMPAR / NMDAR ratio data. Two-tailed t-test comparison of baseline (n = 6, N = 4) against the 24 h post stress is shown in figure S3.A. **B.** There were no significant changes in measures of pre-synaptic release probability, coefficient of variability (CV; P = 0.64) and **C.** paired pulse ratio (PPR; P = 0.66) (stress: n=16, N=8; control: n=22, N=15) measured 24 h post stress. Grey traces illustrate the example traces shown for control condition, scaled to match mean current amplitude for stress condition for comparison. The left panels illustrate individual synaptic response sweeps overlaid to highlight sweep-to-sweep variability. **D.** There were no significant differences between stress and control groups in resting membrane potential (P = 0.94), **E.** membrane resistance (Rm; P = 0.40), nor in **F.** cell capacitance (Cm; P = 0.90). For all:,stress n=16; N=8; control: n=22, N=15. *P < 0.05, **P < 0.01, ***P < 0.001. Data are presented as mean± SEM. Sample sizes are indicated on the figures.

**Supplementary 6.**
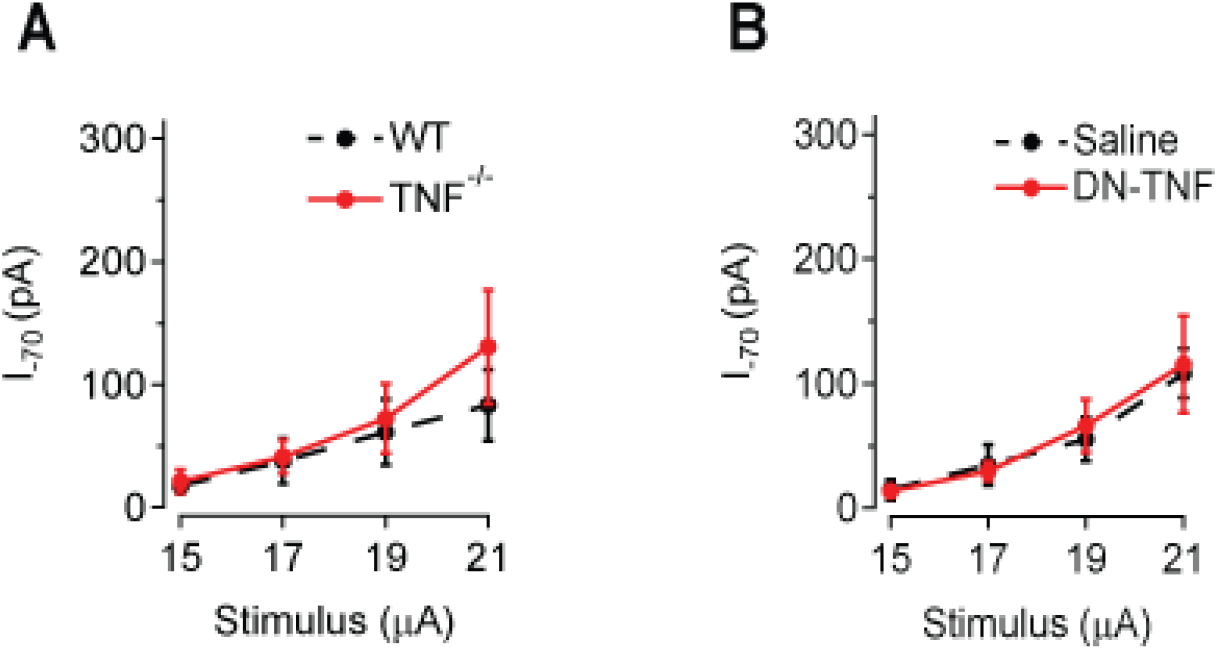
**S6. A.** There was no significant difference in basal AMPAR current in ventral hippocampus SC to CA1 synapses between WT animals and TNF^-/-^ (*P* = 0.16; WT: n=7, N=5; *TNF^-/-^*: n=6, N=3). **B.** Administration of DN-TNF did not alter basal AMPAR current relative to saline-injected controls (*P* = 0.58; Sal: n=18, N=12; DN-TNF: n=17, N=8).

**Supplementary 7.**
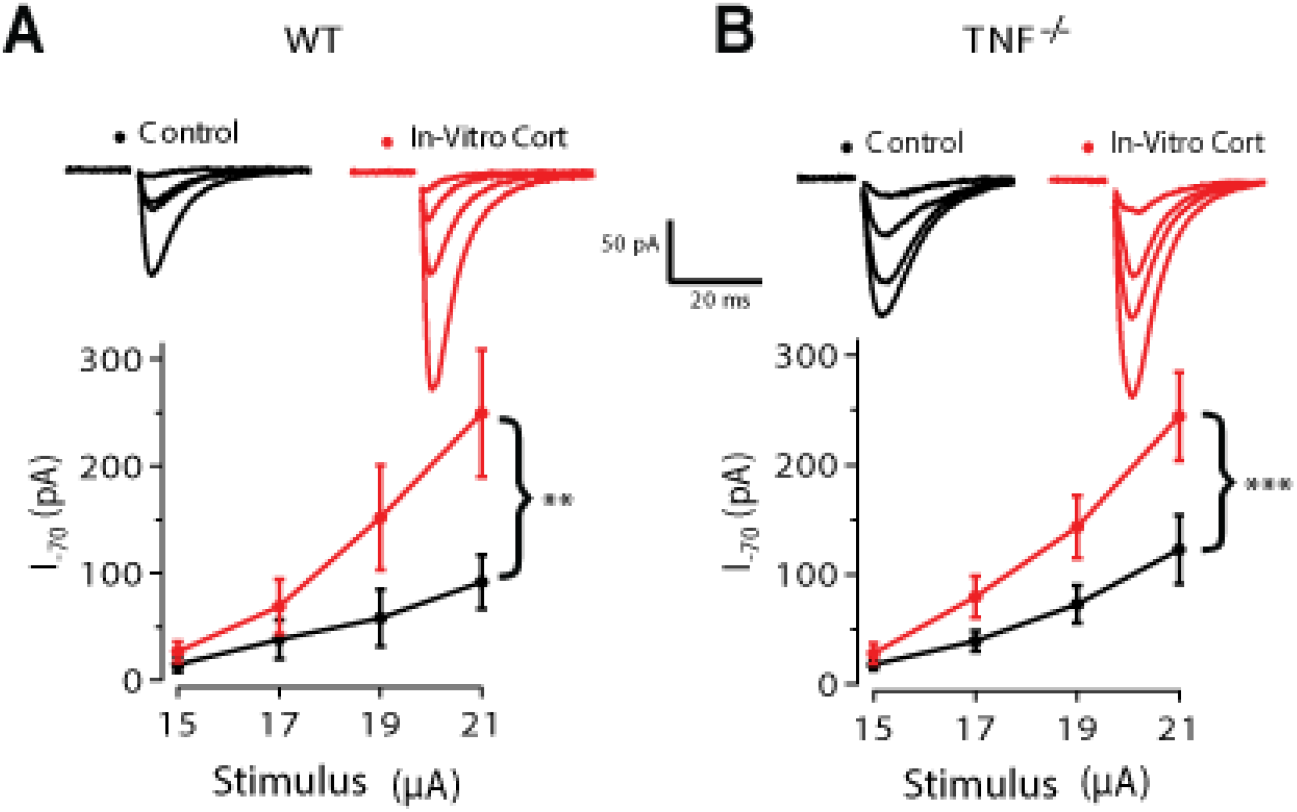
**S7. A.** Ex-vivo application of CORT induces synaptic potentiation of the vHPC-CA1 in both WT (Two-way ANOVA main effect of treatment F(1,34) = 8.3866, P = 0.0066; control: n=4, N = 4; CORT: n = 7, N = 4) and **B.** TNF^-/-^ (Two-way ANOVA main effect of treatment F(1, 48) = 13.8841, P = 0.0005; control: n = 7, N = 5; CORT: n = 7, N = 5). *P < 0.05, **P < 0.01, ***P < 0.001. Data are presented as mean± SEM. Sample sizes are indicated on the captions.

**Supplementary 8.**
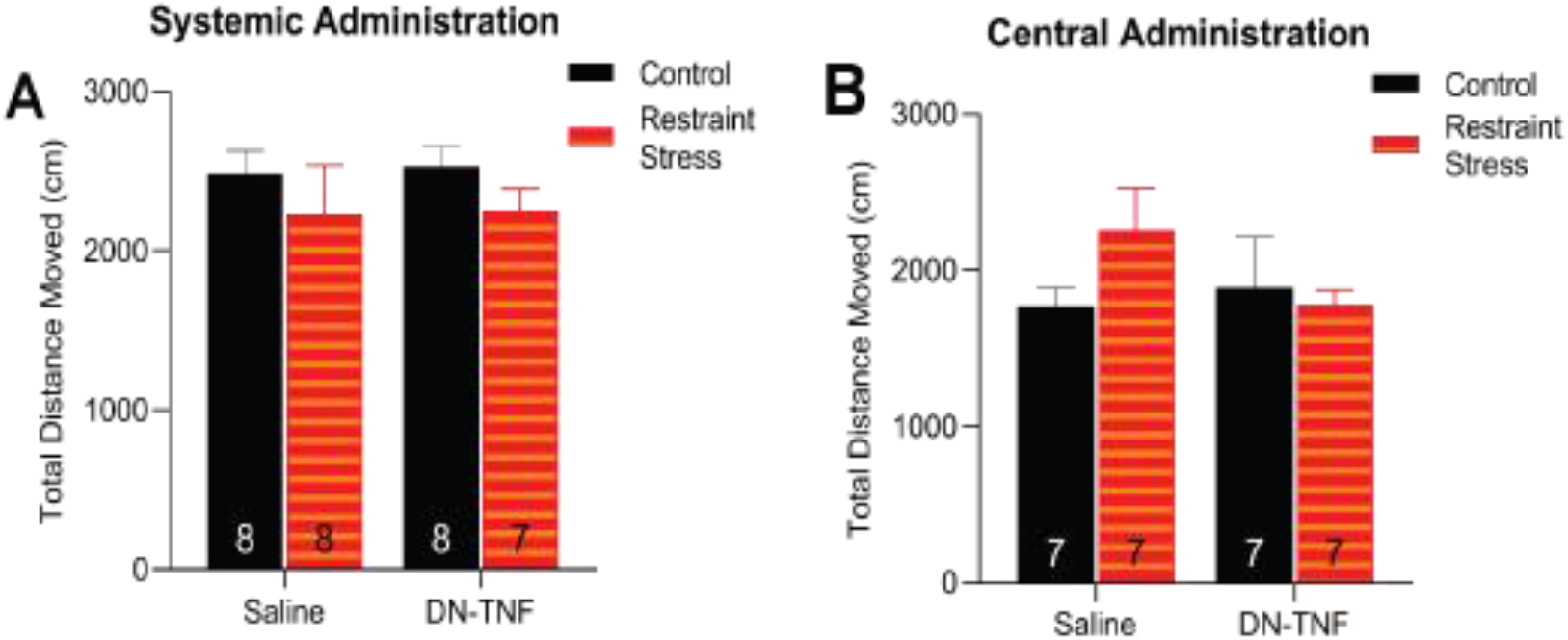
**S8. A.** The observed phenotype is not driven by the effects of the drug treatment on locomotion, as there was no difference in the distance travelled across the different groups (two-way ANOVA of drug treatment F (1, 27) = 0.02533, P = 0.8747; Tukey’s post hoc analyses of control saline versus RS saline P = 0.7978, control DN-TNF versus RS DN-TNF P = 0.7591). **B.** There is no difference in distance travelled under low anxiogenic conditions between the four groups, therefore, the observed behavioral effect of drug treatment is not mediated by differences in locomotive behavior (two-way ANOVA of the main effect of drug F (1, 24) = 0.6220, p =0.4380; Tukey’s post hoc analyses of control saline versus RS saline P = 0.4359, control DN-TNF versus RS DN-TNF P = 0.9860).

**Supplementary 9.**
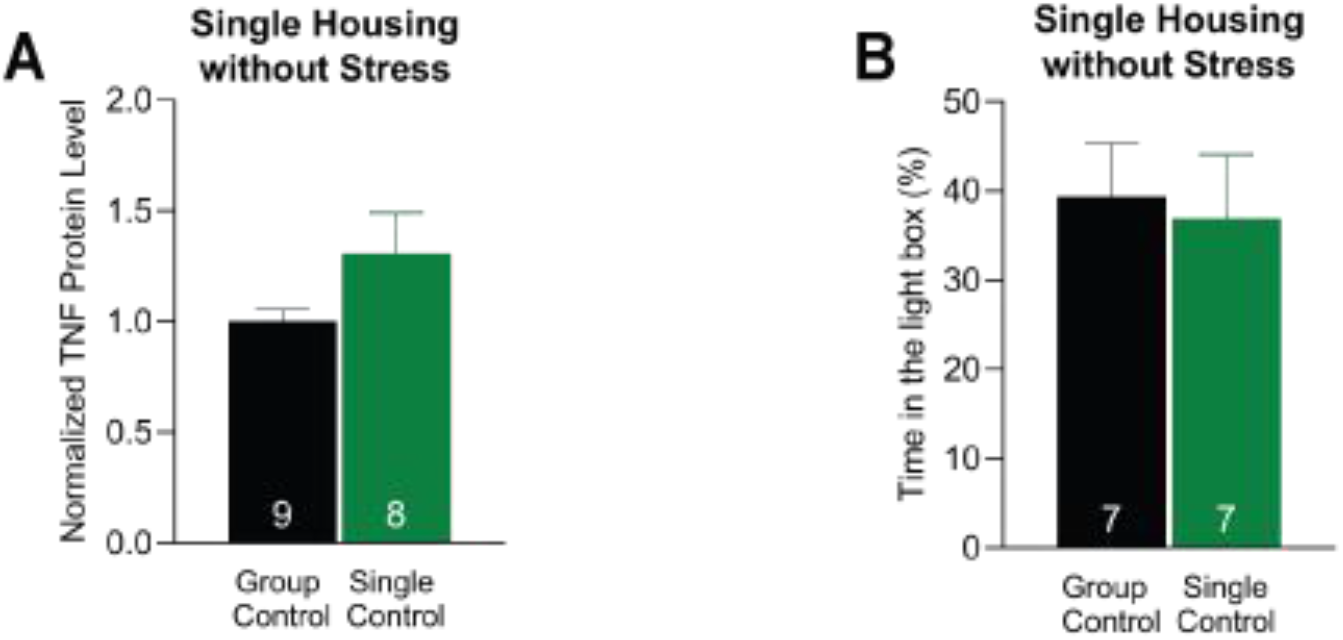
**S9. A.** There is no induction of vHPC levels of TNF in single-housed animals without stress (two-tailed student t-test, P = 0.1124). **B.** Single housing adult WT males over for 24 h does not induce anxiety-like behavior (two-tailed student t-test, P = 0.7999).

**Supplementary 10.**
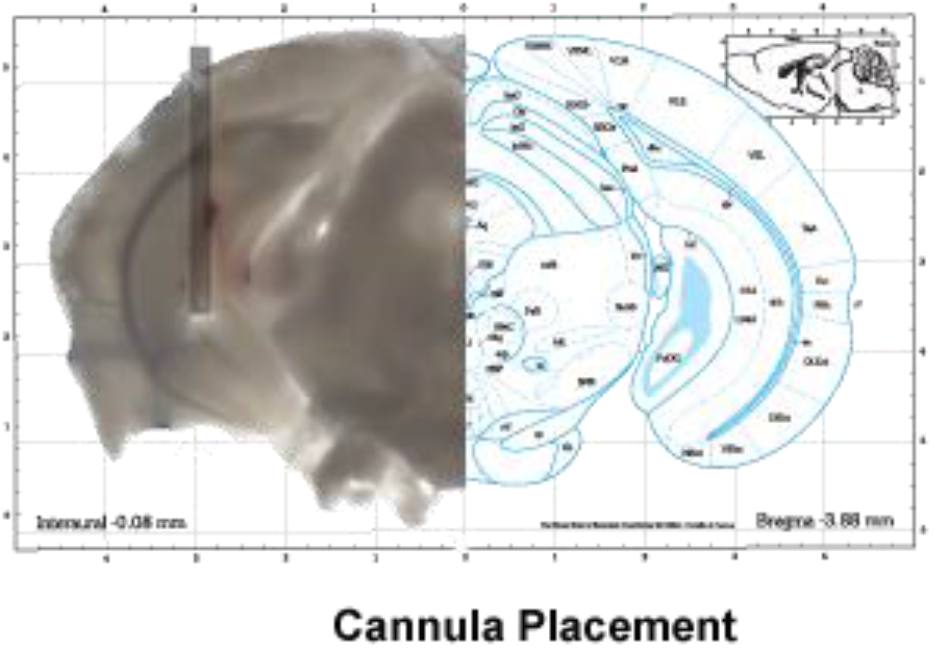
**S10.** A schematic and an image of the placement of the central cannula in the vHPC-CA1. The schematic portion of the figure was obtained from the Mouse Brain Atlas with permission. *P < 0.05, **P < 0.01, ***P < 0.001. Data are presented as mean± SEM. RS, restraint stress. DN-TNF, dominant negative tumor necrosis factor; vHPC, ventral hippocampus. Sample sizes are indicated on the figures.

